# Taste Papillae-associated Salivary Gland Ducts Contribute to Immune Surveillance

**DOI:** 10.64898/2026.01.14.699476

**Authors:** Abdul Hamid Siddiqui, Salin Raj Palayyan, Palmer Wright, Sunil Kumar Sukumaran

**Author notes:** Equal contribution (Abdul Hamid Siddiqui, Salin Raj Palayyan). **Author Contributions:** AHS, SRP and PW: design of the study, performing experiments, acquiring the data, data analysis, interpretation of the results, creation of the figures, and revision of the manuscript. SKS: conceptualization and design of the study, supervision of the data acquisition, interpretation of results, writing of the manuscript. **Competing Interest Statement:** The authors declare no competing interests.

## Abstract

The lingual surface is composed of a tough epithelium that affords effective protection from the resident and food-borne microbiota. The circumvallate taste papillae (CVP) in the back of the tongue are associated with underlying minor salivary glands called von Ebner’s gland (VEG) that drains its secretions into the trenches of the papillae. The taste buds are exposed to the lingual surface through taste pores, to enable interaction with tastants. These openings may be exploited by pathogens for infection. Mucosa-associated lymphoid tissue (MALT) such as those in the gut epithelium, tonsils and lachrymal and salivary ducts possess specialized cells called microfold (M) cells that transport luminal microbes and present them to underlying immune cells, which then initiate an appropriate immune response. Our previous work indicated that type II taste cells might mediate similar immune surveillance akin to M cells and that CVP is patrolled by large population of immune cells. Using single cell RNASeq (scRNASeq) of CVP, we discovered that the duct cells of the VEG strongly express markers for M cells. The ducts were capable of transcytosing fluorescently labelled E. coli and fluorescent nanobeads, which are then taken up by underlying immune cells. We propose that taste and duct cell-mediated immune surveillance might be crucial for preventing infection of taste papillae and for maintaining taste function, and conversely, this pathway might be hijacked by microbes for infection.

## Introduction

The oral microbiome is second only to the intestinal microbiome in complexity and abundance, and the oral cavity is exposed to food and breath-borne microbes as well.(1) The tongue, especially in its dorsal surface, is enveloped by highly keratinized stratified squamous epithelium that affords strong protection from invading microbes. However, taste cells in taste buds extend their microvilli to the luminal surface through taste pores, where they interact with taste molecules.(2) The ducts of the minor salivary glands in the tongue open to lumen as well. These unique anatomical features might represent ‘chinks in the armor’, which microbes might exploit for infection. Muocal tissues such as tonsils, Peyer’s patches, ducts of the salivary, mammary and tear glands etc. contain specialized secondary lymphoid tissue-MALT, harboring M cells that transcytoses luminal microbes across the epithelium and present them intact to immune cells in the underlying lymphoid follicles.(3, 4) These cells process these antigens and generate a tolerogenic or defensive immune response as appropriate. We recently showed that type II taste cells express markers for M cells and might mediate immune surveillance.(5) In the posterior tongue, the CVP is closely associated with the VEG, a serous minor salivary gland that drains its secretions exclusively into the trenches of the CVP through dedicated ducts.(6) Even though mice lack tonsils, their taste papillae and tongue in general are patrolled by a large and diverse set of immune cells.(5, 7) Whether VEG ducts are functional MALT has not been determined so far.

## Results

Droplet-based scRNASeq was used to characterize cellular diversity in the CVP, identifying all known and a few novel (e.g., immature type II cells) taste cell types, non-taste epithelial cells, fibroblasts and immune cells (Figure 1 A). A unique set of epithelial cells were identified as duct cells based on the expression of ductal markers such as *Dcpp1* and *Sox9* and ductal mucins such as *Muc13*. They were classified into ductal and ductal/serous clusters based on a recent scRNASeq study (Figure 1 B).(6) Interestingly, the ductal clusters, particularly the serous/ductal cluster, strongly express M cell marker genes including the receptors for the cytokine RANKL that initiates M cell differentiation *Tnfrsf11a* and *Tnfrsf11b*, the key downstream signaling molecule *Map3k14*, the key transcription factor *Sox8*, and its targets, *Tnfaip2, Gp2* and *Ccl9* (mature M cell marker genes) (Figure 1 B).(3) ). In addition, gene ontology (GO) analysis identified multiple immune function related GO terms enriched in the ductal cells, including those related to canonical and non-canonical (key for M cell regeneration) NF-kappaB signal transduction, leukocyte migration, innate immune response, pattern recognition receptor signaling, defense response to virus etc (Figure 1C). Duct cell clusters also have the highest module score for microfold cell pathways (Figure 1D). The expression of the key M cell transcription factor *Spib* was not observed in the ducts, unlike in type II cells as we reported before.(5) However, it could be strongly stimulated upon RANKL treatment (Figure 1 E & E’). The ducts are surrounded by numerous CD45-expressing immune cells, many of which were attached to duct cells and extended processes in between them (Figure 1 F, F’). We have not determined the identity of these cells, but they had morphological features of dendritic/Langerhans cells and macrophages, which are classical MHC class II-mediated antigen presenting cells. Next, we tested if duct cells could transport luminal microparticles across the epithelium into the mesenchyme by administering fluorescent nanobeads or live mCherry expressing E. coli to the lingual surface.

**Figure 1.**
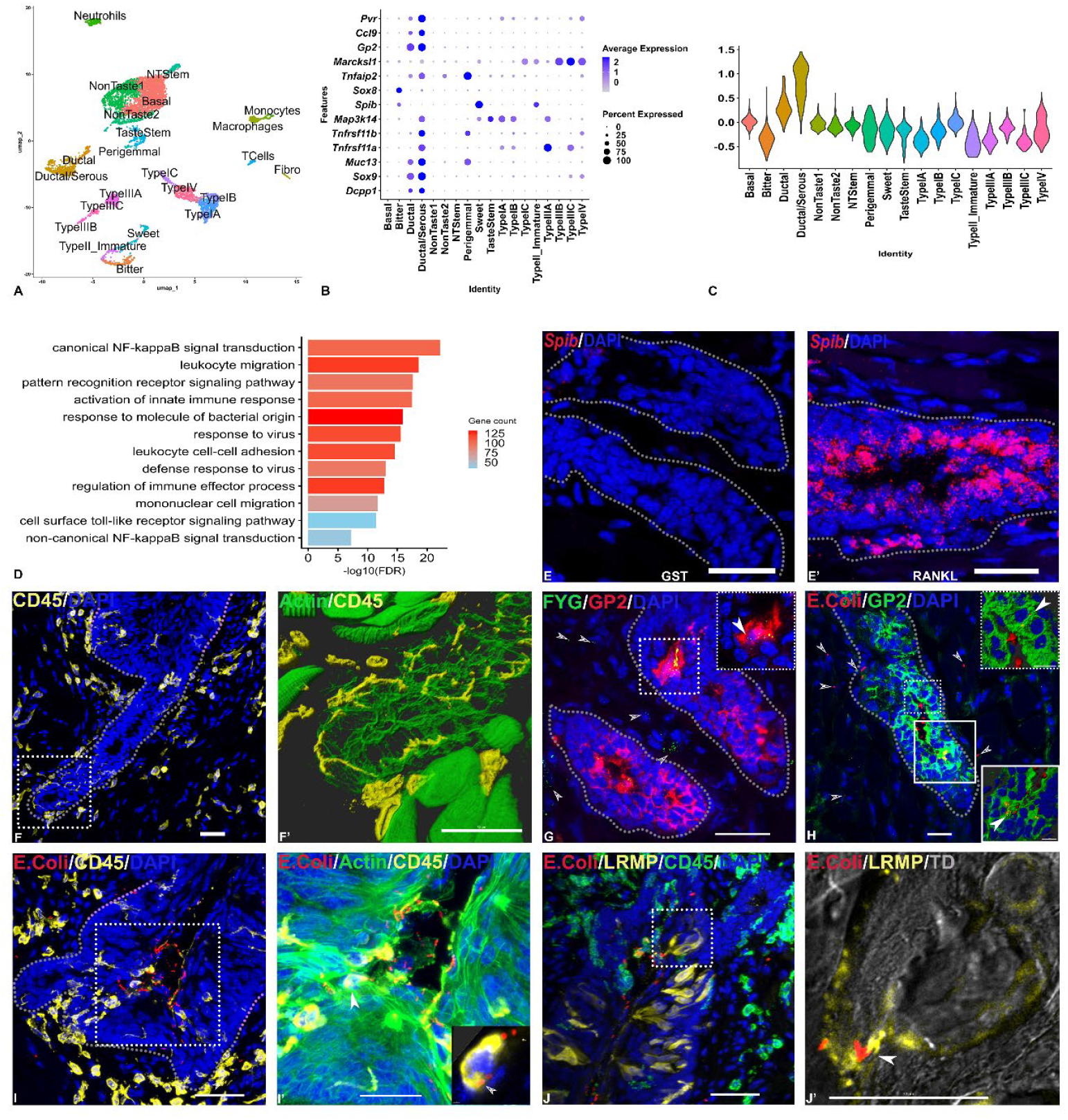
**A)** scRNASeq analysis identifies cell types present in CVP. Two types of ductal cells (Ductal/serous and ductal) were identified in our analysis. **B)** The ductal cells, particularly the ductal/serous cells strongly express M cell marker genes. **C)** The ductal/serous cells had the highest module score for M cell pathway genes. **D)** Ductal cells also were enriched in immune related GO terms. **E, E’)** The key M cell transcription factor *Spib* is not expressed in duct cells in the basal state but could be strongly induced by RANKL treatment. **F)** The VEG ducts are closely associated with multiple immune cells**F’)** Boxed region in F magnified, showing immune cells extending many processes into and between duct cells. **G-H)** Fluorescent nanobeads (G) and E. coli (H) are taken up and transcytosed by GP2-expressing duct cells. Insets show a cluster of duct cells that have internalized nanobeads/E. coli (filled arrowheads). Transcytosed nanobeads/E. coli are highlighted by open arrowheads. **I-I’)** Immune cells were observed invading ducts in CVP in animals administered with E. coli (but not nanobeads). Inset in **I’** is a single slice from a confocal stack showing a CD45-expressing immune cell containing ingested bacteria (arrowhead). **J-J’)**. An E. coli cell is seen bound to the microvillus of a LRMP-expressing type II taste cell. In **J’** a single slice from **J** is shown merged with transmission image (TD) to visualize taste pore and taste bud boundary. In **F’** and **I’**, cell morphology is visualized by staining for actin filaments using phalloidin. In panels **E-J’**, nuclei are visualized by staining with DAPI. Scale bars= 30 µm.

The plasma membrane-anchored isoform of GP2 binds specifically to fimH protein in bacterial fimbriae and nonspecifically to nanobeads and mediates their transcytosis.(3) Nanobeads and E. coli could be seen associated with GP2-expressing duct cells, and some could be observed within the cytoplasm of these cells, presumably undergoing transcytosis (Figure1 G-H). Some nanobeads and E. coli were observed in the underlying mesenchyme, presumably having been trancytosed across the ductal epithelium. We also observed immune cells that have migrated across the ductal epithelium, and ingested E. coli (Figure 1 I). Some E. coli could be seen within taste buds, some of which were inside taste pores, supporting our hypothesis that taste pores are another vulnerable part of taste papillae susceptible to infection (Figure 1 J-J’).

## Discussion

Taste papillae are exposed to microbes and may be susceptible to infection. Salivary glands and epithelial cells secrete many antibacterial substances including immunoglobulin A, lysozyme, defensins, histatins, lactoferrin etc., that limit microbial growth and colonization.(8) The salivary ducts are known to participate in immune surveillance and express many M cell marker genes.(4) We showed that the ducts of the VEG too express M cell marker genes and are capable of microbial transcytosis. They are also intimately associated with immune cells, reminiscent of M cells in well-known MALT. VEG ducts were recently shown to express a host of viral receptors for influenza, HCoV-229E and MERS-CoV45, though not functionally tested.(9) We also detected the expression of poliovirus receptor *Pvr* in ducts (Figure 1B). The taste buds in CVP and foliate papillae (FOP) are located within their trenches and might not get salivary supply from the main salivary glands. This might have necessitated their close association with VEG. CVP and VEG develop from the same anlagen during embryonic development and remain associated with each other in adulthood as well. In addition, *Lgr5*-expressing stem cells in the ducts contribute to the regeneration of both taste and non-taste epithelial cells in the CVP and the VEG.(6) Thus, the unique relationship between CVP (and presumably FOP) and VEG appears to be essential for regeneration, taste sensation and immune surveillance. Duct cells might be exploited by microbes that can hijack the transcytosis pathway for local and systemic infection.

## Materials and Methods

### Animals

Two-three-month-old C57BL/6J mice were used for all experiments. Animal experiments were performed in accordance with the National Institutes of Health guidelines for the care and use of animals in research and reviewed and approved by the Institutional Animal Care and Use Committee at University of Nebraska-Lincoln (protocol # 2610 and 2366 to SKS). Animals were housed in a specific pathogen-free vivarium with a 12-h light/dark cycle and open access to food and water.

### Fluorescent beads and Bacterial transcytosis experiments using mCherry expressing E. coli

E. coli BL21DE3/DH5α strains were transformed with pON mCherry plasmid (kind gift from Jennifer M Auchtung, Department of Food Science, UNL). A single colony of transformed bacteria harboring expression plasmids were selected and grown in Luria-Bertani (LB) media supplemented with 25 µg/ml chloramphenicol and grown to late log phase. They were harvested by centrifugation and resuspended in sterile phosphate-buffered saline (PBS) to a final volume of 1.0 ml. Mice were anesthetized by intraperitoneal injection of ketamine (4.28 mg/ml), xylazine (0.86 mg/ml), and acepromazine (0.14 mg/ml) at a dosage of 10 ml/kg. Following induction of anesthesia, mice were tracheotomized as described previously.(10) The bacterial suspension or fluorescent polystyrene latex nanobeads (Fluoresbrite YG; 200 nm diameter; Cat. no. 09834-10; Polysciences, Warrington, PA) was administered orally and incubated for 60 minutes. At the end of incubation, animals were transcardially perfused with 4% PFA. The tongue was removed, washed thoroughy to remove adhered bacteria. It was then fixed in 4% PFA for 1 hour and cryoprotected by overnight immersion in 30% (w/v) sucrose. Tissue was embedded and frozen in optical cutting temperature (OCT), sectioned (10/12 µm) using a cryostat and used for immunohistochemistry as described below.

### RANKL preparation and injection

The expression and purification of recombinant mouse RANKL were performed as previously described with some modifications.(11) The missing 5’ 127 nucleotides of RANKL sequence was synthesized using gBlock gene fragment (IDT) and then cloned into pGEX-4T-2 RANK-L vector (Plasmid no. #108571; Addgene Watertown, MA) using NEBuilder HiFi DNA Assembly Master Mix (New England Biolabs, Ipswich, MA). The insert was verified by sequencing and then transformed into *E. coli* strain for expression. Bacteria harboring expression plasmids were selected and grown in Luria-Bertani (LB) media supplemented with 100 µg/ml ampicilin. The cultures were induced with 40 μM isopropyl -D-1-thiogalactopyranoside (IPTG) for 16 h at 25°C, and the glutathione-S transferase (GST) tagged RANKL (GST-RANKL) were purified from bacterial lysate by affinity chromatography on Pierce Glutathione Spin Columns (Cat. no. 16108, Thermo Scientific, Waltham, MA) followed by dialysis against multiple changes of phosphate buffered saline (PBS), pH 7.4. Recombinant GST used as a control was prepared by the same method using empty pGEX-5X-2. GST or GST-RANKL, with at least 95% purity as demonstrated by SDS-PAGE was administered to WT C57BL/6J mice age between 4-6 weeks by intraperitoneal injections *(250* μg/day/mice for three days). Mice were sacrificed at day 4 and lingual epithelium or lingual tissue were collected for histology.

### Immunohistochemistry

Standard immunohistochemical techniques were used as previously described.(11) Briefly, frozen sections were rehydrated with PBS. Nonspecific binding was blocked with SuperBlock Blocking Buffer (Cat. no. 37580, Thermo scientific, Waltham, MA) at room temperature for 1 h. Sections were incubated with primary antibodies [CD45 (1:500; Cat. no. AF114, R and D systems, Minneapolis, MN), GP2 (1:500; Cat. no. D278-3; MBL Life sciences, Japan), and LRMP (1:500; Cat. no. ORB166443; Biorbyt, Durham, NC)] diluted in blocking buffer for overnight at 4 °C in a humidified chamber. After three 15-min washes with PBST, slides were incubated for 1 h at room temperature with one of the secondary antibodies (1:500; Cat no. 705-606-147, 712-545-150, 112-545-008; Jackson Immuno Research, West Grove, PA) in blocking buffer. F-Actin staining was done using phalloidin (1:100; Cat. no. R37110, ActinGreen 488 ReadyProbes reagent; Thermo Fisher Scientific, Waltham, MA) for 30 mins followed with DAPI (1:1000; Cat. no. D1306; Invitrogen™, Thermo scientific, Waltham, MA) to label cell nuclei for cell counting.

### RNAScope Hiplex assay

RNAscope assay was done using the Hiplex fluorescent assay kit for mice (Advanced Cell Diagnostics, Hayward, CA, Cat. no. 324443) with RNAscope™HiPlex Probe-Mm-Spib-T3 (Cat. no. 408781-T3) as previously described using the manufacturer’s instructions.(11) Positive and negative control probes were run in parallel to test probes to ensure proper hybridization and imaging conditions were attained in our experiments. Confocal images were captured using a Nikon A1R-Ti2 confocal laser scanning microscope using NIS-Elements A1R software image acquisition and analysis software, using 40/60X objectives. Images were taken using a sequential channel series setting to minimize cross-channel signal, and the channels used were 408 nm for dapi and 650 nm for Spib. Z-series stack with 10 images per stack was captured at a step size of 1 μm. Acquisition parameters [i.e., gain, offset, photomultiplier tube (PMT) setting] were held constant between experiments.

### scRNASeq of taste papillae

Chromium™ Single Cell 3′ Solution (10x Genomics Inc, Pleasanton, CA, Cat. no. PN-1000268) was used for scRNASeq analysis. Single cell preparation from CVP was done as previously described.(12, 13) A protease cocktail was injected under the lingual epithelium of excised tongue (n=16 mice) and incubated at 37°C for ten minutes. The epithelia were peeled and the CVP were excised and minced to form single cells. Single cell capture, library preparation, sequencing, and primary analyses of sequencing data were done using 10X genomics protocols, and secondary analysis was done using the Seurat package in R.(14, 15) Ambient RNA and doublets were removed using SoupX and Doubletfinder packages in R. (16, 17) The codes for scRNASeq secondary analysis in seurat will be uploaded to the lab’s github page.

## Acknowledgments

We wish to thank Brian Lewandowski for help with scRNASeq and Jennifer Auchtung for the mCherry expressing E. coli strain. This work was supported by National Science Foundation CAREER award # 2443659, NIH/NIDCR New Investigator RO3 award #1R03DE032417, a Project leader award from Nebraska Center for the Prevention of Obesity Diseases (NIH-NIGMS grant no. P20GM104320), and PA State tobacco grant STA019A01SUKUM to SKS.

